# Brain-specific PP1/LRRK2 interaction-targeting peptides as a novel strategy for Parkinson’s disease treatment

**DOI:** 10.64898/2026.07.27.740889

**Authors:** Severine Loisel, Fanny Petit, Gwenaelle Auregan, Martine Guillermier, Marjorie Benfissa, Pauline Gipchtein, Asma Touffeil Mohamed, Kristina Posnograjeva, Tambet Teesalu, Romina Aron Badin, Angelita Rebollo

## Abstract

The LRRK2 kinase has emerged as a priority therapeutic target due to its well-documented role in Parkinson-s diseases (PD). One of LRRK2’s key partners is the phosphatase PP1, which dephosphorylates LRRK2. Therefore, modulating this protein/protein interaction represents a promising therapeutic strategy for PD.

We have developed a bi-functional peptide, PEP 3, capable of crossing the blood-brain barrier (BBB) and disrupting the LRRK2/PP1 interaction. *In vitro* competition assays confirmed that PEP 3 specifically targets this interaction. The *in vivo* imaging demonstrated that the peptide remains detectable in the mouse brain up to 6 hours or retro-orbital vein post-injection. The PEP 3 peptide does not show chronic toxicity in CD1 mice and is resistant to degradation by mouse, human, dog and monkey serum proteases. In mouse models overexpressing alpha-synuclein, repeated intraperitoneal injections of PEP 3 for 15 and 30 days were well tolerated. Immunohistochemistry revealed a significant reduction of dopaminergic cell death in the substantia nigra, and quantification of phosphorylated pathological alpha-synuclein showed decreased levels in the PEP 3-treated group compared to controls. These findings support the potential of PEP 3 as a therapeutic agent for Parkinson’s disease.

## INTRODUCTION

Neurological disorders are the most common cause of human disability. Among these, Parkinson’s disease (PD) shows an increasing incidence in the general population over 60 years old (***1***). Clinically, PD is characterized by motor symptoms like bradykinesia, often accompanied by resting tremors, rigidity or postural instability as well as non-motor symptoms including sleep disturbance, executive disfunction and gastrointestinal problems. The two main pathological hallmarks of PD are the selective loss of dopaminergic neurons of the substantia nigra (SN) that can reach 50% by the time of diagnosis, (***2***) and the accumulation of aggregated α-synuclein and other proteins in Lewy bodies (***3, 4***).

The increasing prevalence of PD and its profound impact on the quality of life of patients underscore the urgent need for developing treatments that slow or halt disease progression. Advances in our understanding of PD pathogenesis have identified of several potential therapeutic targets for disease treatment. However, to date, no treatment has demonstrated disease-modifying effects on PD (***1***).

One major challenge in developing effective PD therapies is the delivery and biodistribution of therapeutics. The blood-brain-barrier (BBB) restricts the passage of molecules into the brain, protecting it from potentially harmful from blood-borne substances (***5***). Exploiting the natural transport mechanisms of the BBB, that can be broadly classified as passive or active, depending on energy requirements, offers a promising strategy for brain drug delivery. Inspired by existing proteins in the brain, peptide shuttles can harness these mechanisms to facilitate the transport of compounds that cannot naturally cross the BBB (***6***). For example, peptides derived from or targeting apolipoproteins have been extensively explored. In addition, phage display is also a rich source of bioactive BBB shuttle-peptide candidates (***7***). Using *in vivo* peptide phage biopanning, we recently developed a potent brain-targeting and - penetrating peptide capable of delivering a variety of cargoes, ranging from small molecular payloads to nanoparticles. Following *in vivo* tests in mice and rats, as well as *ex vivo* tests on human samples, the peptide demonstrates efficient uptake across multiple brain regions following systemic administration, highlighting its potential for treating a wide range of neurological and neuro-oncological diseases.

Protein-protein interactions (PPIs) are recognized as promising therapeutic targets. Interfering peptides (IPs) which can disrupt PPIs are gaining attention because their physicochemical properties make them better suited than small molecules to interfere with the large surfaces involved in PPIs. Advances in peptide administration, stability, biodistribution, cost effectiveness and safety have further fuelled the interest in peptide-based drug development. Following key *in vitro* and *in vivo* proof of concept experiments, several IPs targeting have been preclinically validated and some are already under clinical development for cancer (***8–11***). IPs combine selective and specific targeting, with a good efficacy and potential for intracellular targeting. Moreover, they have a predictable metabolism and very low toxicity, and their intracellular concentration is not dependent of flow pumps (as is the case for small molecules), thereby bypassing efflux-based resistance mechanism. Finally, they are much less immunogenic than recombinant proteins or monoclonal antibodies. Taken together, IPs are excellent candidates to become new therapeutics.

Among potential PPI targets, serine/threonine phosphatase 1 (PP1) and phosphatase 2A (PP2A) are the most widely distributed and abundant serine/threonine phosphatases in eukaryotic cells since they regulate essential cellular functions such as proliferation, apoptosis, and memory. (***12, 13***). In vertebrates, about 200 proteins have been described as partners or interacting with PP1. Many PP1 partners have different domains for their association with PP1, for substrate recruitment, and for subcellular targeting. It is thought that this diversity in binding modes underlies the high specificity of PP1 functions *in vivo* (***14, 15***). Leucine-Rich Repeat Kinase 2 (LRRK2) is a PP1-interacting protein (***16***). Mutations in the LRRK2 gene cause familial PD and constitute a significant risk factor for sporadic PD. A cluster of phosphorylation sites within LRRK2 (***17–20***) contributes to PD pathology, as several PD-linked LRRK2 mutations have been associated with a dephosphorylated state. Notably, LRRK2 is dephosphorylated following inhibition of its kinase activity, an effect that has been proposed as a therapeutic approach for PD (***21, 22***).

Building on our previous success using IPs to target PPIs, we report here the preclinical validation of a bi-functional peptide capable of crossing the BBB and disrupting the interaction between the phosphatase PP1 and LRRK2, positioning it as a promising therapeutic candidate for the treatment of PD.

## MATERIALS AND METHODS

### Peptides synthesis and sequences

The sequences of the peptides used in this study are shown in Table I. The sequence of the IP blocking the interaction PP1/LRRK2 is GFWSRLINRLLEI. This peptide was combined with BBB homing/penetrating modules via linkers — aminohexanoic acid (Ahx) or the tripeptide GGS — to provide structural flexibility. Peptide PEP 1 and PDP2 were used as positive controls. The BBB shuttle sequence of these peptides corresponds to the already published Angiopep (sequence: TFFYGGSRGKRNNFKTEEY). In addition, the novel peptide BBB shuttles used in this study were: Cereper (sequence: SRRVISRAKLAAAL) and CVG (https://worldwide.espacenet.com/patent/search?q=pn%3DEP4577229A1). Peptides Lin TT1-IP and BBB3 were used as negative and positive controls for BBB penetration respectively. Peptides were synthetized in an automated multiple peptide synthesizer with solid phase procedure and standard Fmoc chemistry (GL Biochem (Shanghai, China). The purity and composition of the peptides were confirmed by reverse phase high performance liquid chromatography and by mass spectrometry (MS).

**Table I.**
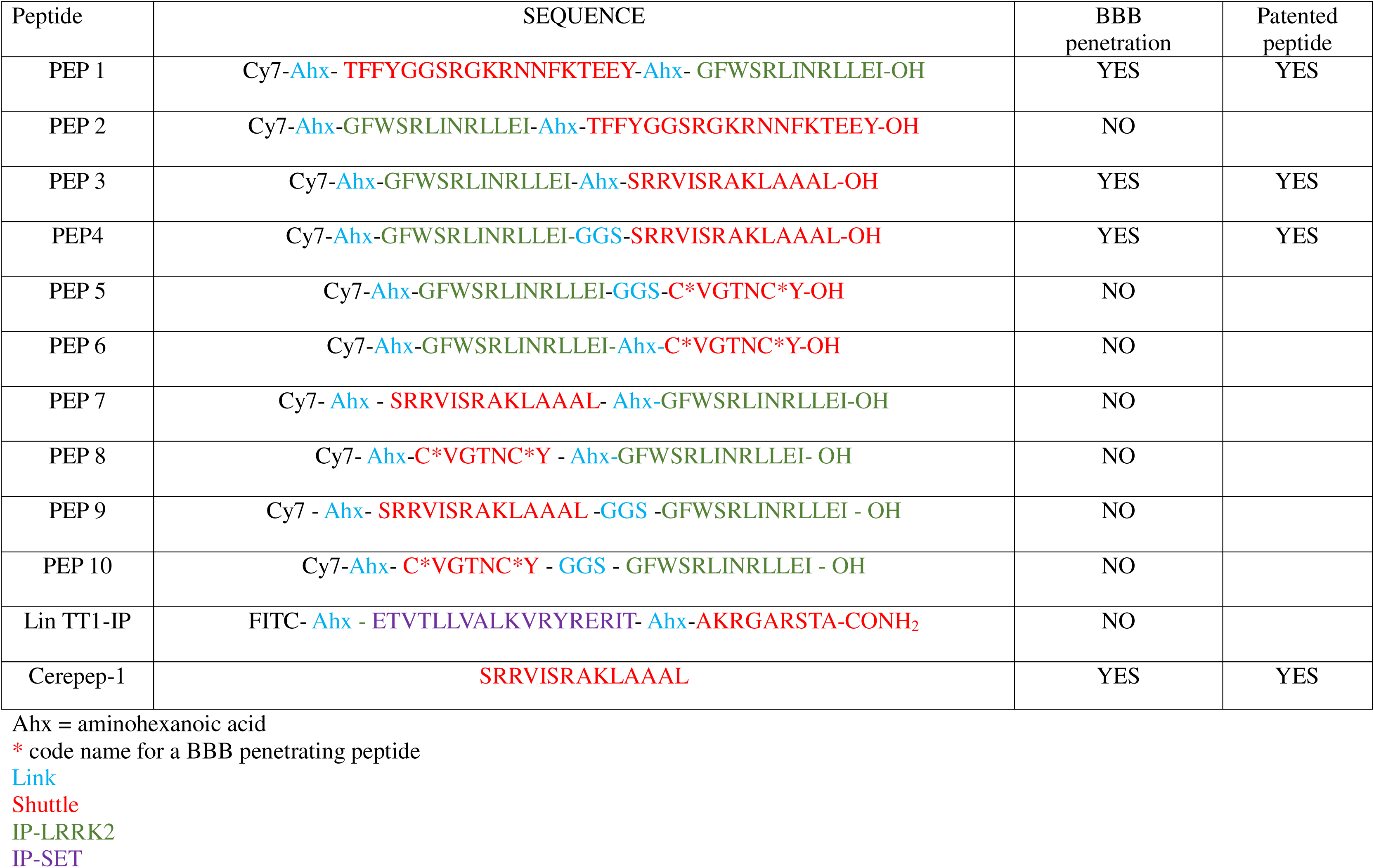
Sequence and structure of the peptides generated and used in this study. The colour code shows the different parts of the peptide. Cy7, sulfo-cyanine 7, is the fluorescent dye. The linker is shown in light blue (Ahx = aminohexanoic acid). The sequence of the shuttle appears in red and the code name for a BBB penetrating peptide is marked by an asterisk (*). The sequence of the IP-LRRK2 peptide is shown in green and that of the control IP-SET in purple.

### Cell line

The human cell line MDA-MB-231 (ATCC HTB-26) was cultured in Dulbecco’s Modified Eagle Medium (DMEM), medium supplemented with 10% foetal call serum (FCS, Thermo Fischer).

### In vitro PP1/LRRK2 interaction competition

The protocol for isolation of lysates from the MDA-MB-231 cell line was previously described (***23***). Briefly, 5×10^6^ cells were lysated for 20 min at 4°C in lysis buffer (50 mM Tris HCl pH 8, 1% NP40, 137 mM NaCl, 1 mM MgCl_2_, 1 mM ClCa_2_, 10% glycerol) and protease inhibitor mixture. The protocol for isolation of proteins from brain tissue was also previously described (***24***). Briefly, brain tissue of CD1 mice was cut into small pieces and transferred to a tube of Precellys beads CK28-R. A total of 400 μl of lysis buffer was added (Tris 50 mM pH8, 150 mM NaCl, 1% Triton), and supplemented with protease inhibitors. The tube was placed in the Precellys Evolution machine and shaken for 30 s followed by a 30 s pause, and this cycle was repeated 5 times. The extracts were centrifuged and the supernatant was transferred to a fresh tube and centrifuged at 12,000 rpm for 20 min at 4^°^C. The supernatant was recovered to estimate the protein concentration.

Both, cell line and brain tissue lysates (500 μg) were immunoprecipitated with anti-PP1 antibody overnight at 4^°^C and protein A/G Sepharose was added for 1h at 4^°^C. After several washing steps in 1x TBS (20 mM Tris HCl pH7.5, 150 mM NaCl, 0.05% Tween 20), a competition assay using 1 mM of the IP (GFWSRLINRLLEI), the shuttle alone or an irrelevant antibody for 30 min at room temperature (RT) was performed 4 times to interfere with the PP1/LRRK2 interaction. Following additional washing steps, immunoprecipitates were separated by SDS-PAGE, transferred to nitrocellulose membranes, and blotted with anti-LRRK2 antibody. Membranes were then washed and incubated with PO-conjugated secondary antibody. Protein detection was performed using ECL system. As an internal control, the membranes were also blotted with an anti-PP1 antibody. Densitometric analysis of the blot was performed using ImageJ software.

### Characterization of PP1/LRRK2 peptide interaction by ELISA

A total of 100 μM of biotinylated IP-LRRK2 peptide diluted in 100 μl of PBS was incubated for 2h at RT in a 96 well-streptavidin coated plate. Wells were washed with PBS/0.5% Tween 20 (PBST) and loaded with 100 μl of PP1 in a 2.5% BSA-PBS solution for a final concentration of 2 μM. The plate was incubated over night at 4^°^C and washed with PBST. Anti-PP1 antibody was added for 1h at RT. Wells were washed with PBST and incubated with an HRP-conjugated secondary antibody for 1h at RT. Wells were washed with PBST and TMB substrate was added and incubated for 30 min at RT. The reaction was stopped with 2N sulphuric acid and the absorbance was measured at 450 nm.

### Animals and housing for immunochemical staining and toxicity

A total of 85 4-8 weeks old CD1 mice were purchased from Charles Rivers Laboratories. Mice were housed in ventilated cages in group of 5 under the following standard conditions: water and food “ad libitum”, ambience temperature maintained at 22°C ± 1°C and alternance of light and dark periods of 12h each. All experiments were approved by the Ethics Committee (Fluorescence and body weight:37630_2022061011291086; PD: #45289-2023102514188124) and conducted in accordance with the EU directives.

### Fluorescence live imaging

Two groups of 6 CD-1 mice (female, 6-8 weeks old) were used for live imaging experiments. Before starting the experiment, animals were given local anesthesia by tetracaine 1% eye drops and anesthetize with isoflurane (4% air-isoflurane blend). Animals were then injected into the retro-orbital vein with the Cy7-labelled corresponding peptides listed in Table 1, and placed in the acquisition chamber of an *in vivo* imaging system equipped with a cooled slow-scan CDD camera and driven with Indigo software. Acquisition of fluorescent images was performed at different times after injection and at different peptide concentrations to study biodistribution. Images were captured at 1×1 binning, with exposure times ranging from 0,1 to 1.0 s, depending on the fluorescence intensity.

### Analysis of peptide integrity on human, mouse, dog and monkey serum

The PEP 3 peptide was incubated at 37^°^C with 250 μl of human, mouse, dog or monkey serum for different periods of time ranging from 1 to 24h. The Lin TT1-IP peptide was incubated at 37°C with 250µl of human serum from 1 to 24h. After each incubation time, samples were collected and peptide degradation stopped by freezing. Peptides were extracted from samples using the Proteo-miner Protein Enrichment system. The percentage of intact peptide was estimated by MS using MALDI-TOFF as previously described (***10***). Measurements were performed in triplicate and results were analysed using the Clipro tools, Flex analysis software (Bruker).

### Surgery and treatment administration

An AAV viral vector, produced in HEK932 cells and carrying the mutant human alpha-synuclein (A53T) gene under the control of the PGK neuronal promoter was stereotaxically injected to bilaterally overexpress alpha-synuclein in the substantia nigra (SN) of adult CD-1 mice.

All mice received analgesia (0.05 mg/kg Buprenorphine, sub-cutaneous, s.c.) 15 min before surgery, followed by anesthesia with a mixture of Ketamine (100mg/kg, s.c.) and Domitor (0.25mg/kg, s.c.). Local analgesia was applied at the incision site (5 mg/kg xylocaine, s.c.) and hydration was controlled during the procedure (250µl of warm NaCl 0.9% s.c. on the back). Body temperature was kept constant at 37.5°C by a retrocontrolled thermic mat (RWD VI-69026 Pad). The skin was disinfected and incised, the skull exposed, and a burr hole was drilled at 2 injection coordinates (AP:-2.8mm, L:+/-1.6mm, Z:-4.5mm regarding the skull bone, tooth bar: 0mm). A volume of 1µl of virus at a concentration of 2.5×10^10^ Vg per site was stereotaxically injected at a rate of 0.25µl/min using a CMA 4004 Syringe pump. The wound was sutured and the skin disinfected before reversal of anesthesia (0.25 mg/kg atipamezole, s.c.). One month after viral injection, animals were administered 7.5mg/kg of the PEP3 peptide or PBS intra-peritoneally (i.p.) once a day, 5 days a week, alternating the injection on the right and left side every day. The first cohort of mice (n=20) received the treatment (n=10) or PBS (n=10) for 2 weeks (w), and the second cohort of mice (n=10 PEP3 and n=10 PBS) for 4 w. Clinical follow-up included daily health monitoring and weekly weighing to detect any acute or semi-chronic adverse reactions to the PEP3 treatment.

### Immunohistochemical staining

At the end of the PEP 3 treatment period, animals were given analgesia (0.3 mg/kg buprenorphine, s.c.) 30 min before termination and were deeply anesthetized (330 mg/kg ketamine +1.2 mg/kg domitor, s.c.) before intracardial perfusion (8ml/min) with PBS (0.1M, 1 min) followed by 4% PFA-PBS (10 min). Brains were dissected and post-fixed in PFA 4% for 24h, before immersion in cryoprotectant (30% sucrose-PBS). Brains were then sliced (40µm) in a freezing microtome and incubated with antibodies directed against tyrosine hydroxylase (TH) (1/600; Merck MAB318 (Mouse) - clone LNC1) and a secondary fluorescent antibody (594mn), or phosphorylated alpha-synuclein (P129 a-Syn) (1/1000; Abcam Ab51253 (Rb) - EP1536Y clone) and a biotinylated secondary antibody (1/1000) followed by an avidin-biotin amplification and DAB. All stained slices were scanned in a Z1 AxioScan (Zeiss). The TH results were obtained by measuring fluorescent intensity in the SN using the ZEN software. Alpha-synuclein results were obtained by thresholding positive staining in the region of interest using the FIJI software.

### Statistical analysis

A power calculation was performed on a freeware online tool (https://eda.nc3rs.org.uk/experimental-design-group#PowerCalc) to determine the sample size for the *in vivo* efficacy studies at 2 and 4 w of treatment, assuming a 50% effect size, 30% variability, a 0.05 significance level and 0.9 power.

Data were analyzed using Welch’s t-test to compare the means between the PBS and PEP3 groups, as this method accounts for unequal variances between samples. A one-tailed approach was employed because our hypothesis was directional, specifically predicting a decrease in phosphorylated alpha-synuclein scores and an increase in TH staining in the treated group. Prior to hypothesis testing, the normality of the distributions was assessed using the Shapiro-Wilk test. Although either group displayed a departure from normality under one condition or another, a parametric approach was maintained due to the inherent robustness of the t-test against moderate non-normality. The magnitude of the difference between groups was quantified using Hedges’ g to provide a corrected estimate of the effect size. Statistical significance was defined at p=0.05. All calculations, including the determination of the 95% confidence intervals and the statistic, were performed using R version 4.3.

## RESULTS

### Assessment of the specificity of the interfering peptide IP-LRRK2 and PEP 3 for blocking the PP1/LRRK2 interaction

Using the PEP scan approach, we have identified the binding site of LRRK2 to PP1 (***23***). This sequence, GRWSRLINRLLEI, referred to as IP-LRRK2, was associated to a shuttle (Cerepep 1) able to cross the BBB. Lysates from MDA-MB 231 cells were immunoprecipitated with anti-PP1 antibody and an *in vitro* competition assay was performed using 1 mM of PEP 3 peptide, the shuttle alone or an irrelevant peptide (peptide blocking Ras/Raf interaction). Figure 1A shows that only PEP 3 is able to block PP1/LRRK2 interaction. Total extracts or immunoprecipitates without peptide were used as negative controls (Fig 1A, already published in PlosOne, 2020).

**Figure 1.**
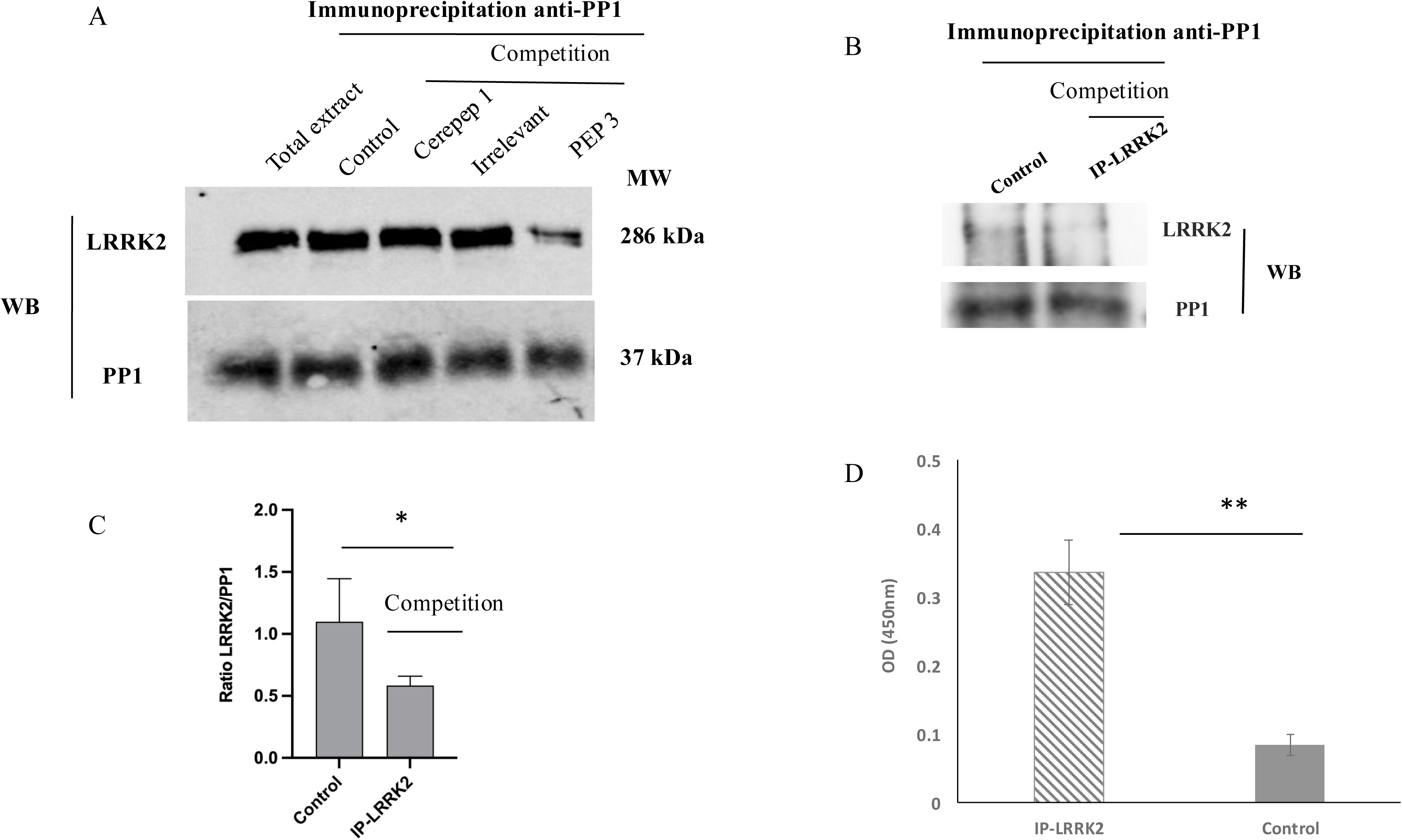
*In vitro* competition of PP1/LRRK2 interaction by IP-LRRK2 and PEP 3 peptides and binding of IP-LRRK2 to PP1 protein. **A)** The interaction PP1/LRRK2 was assessed *in vitro* on immunoprecipitates derived from cell lysates in the presence or absence of the interfering PEP 3 peptide. The competition assay was performed with 1 mM of the shuttle alone (Cerepep 1), an irrelevant peptide (interfering peptide blocking Ras/Raf interaction) or without peptide as controls. PP1 expression was used as internal control of protein loading. This experiment was previously published (*DOI: 10.1371/journal.pone.0237110)* **B)** Brian tissue from CD1 mice was lysed and cytoplasmic extracts immunoprecipitated with anti-PP1c antibody. The PP1/LRRK2 interaction was assessed by *in vitro* competition assay using mouse brain-derived PP1 immunoprecipitates. Immunoprecipitates were blotted with anti-LRRK2 and anti-PP1 antibodies, the latter as an internal control for protein loading. **C)** Densitometric analysis of four independents blots (p< 0.05) (B)**. D)** Direct binding of the biotinylated IP-LRRK2 peptide to PP1 protein was assessed by ELISA. Binding of PP1 is expressed as the mean OD at 450 nm of triplicate wells. Bars indicate standard deviation (SD). Similar results were obtained in two independent experiments (p < 0,05).

In order to further validate that the IP-LRRK2 targets the LRRK2/PP1 interaction, we performed an *in vitro* competition experiment in cell extracts from brain tissue. Brain lysates isolated from adult CD1 mice were immunoprecipitated with anti-PP1 antibody and the interaction with LRRK2 was assessed in the presence or absence of the IP-LRRK2 peptide. LRRK2 was detected in the control anti-PP1 immunoprecipitates whereas its signal was markedly reduced after competition with 1 mM of the IP-LRRK2 (Fig 1B). Precipitated PP1 levels were used as an internal control and were similar under both conditions (Figure 1B). Densitometric analysis of western blot bands of four independent experiments suggested a reduction of the LRRK2/PP1 interaction in the presence of the IP-LRRK2 peptide (Figure 1C).

We next used an ELISA test to show that the biotinylated IP-LRRK2 peptide could directly bind to PP1 protein (Figure 1D). Together, these results confirm the specificity of IP-LRRK2 and PEP 3 peptides for blocking PP1/LRRK2 in different cellular models.

### Bi-functional peptides can cross the BBB

Table I summarize the structure and the sequence of the 10 IP-LRRK2 peptides used in this study that were generated by combining 3 different shuttles able to cross the BBB, and 2 different linkers. Moreover, the IP peptide linked to a tumor homing linTT1 peptide was used as a negative control that does not cross the BBB, while shuttle Cerepep-1 was used as positive control. The linkers employed between the shuttle and IP module were either the tripeptide GGS or the aminohexanoic acid, providing flexibility to the construction. Out of the the BBB shuttle modules used, one was the widely used clinical-stage Angiopep-2 (TFFYGGSRGKRNNFKTEEY) while the other two brain shuttles (Cerepep-1 and CVG, SRRVISRAKLAAAL and C*VGTNC*Y respectively) were generated de novo in Dr Teesalu’s laboratory.

Furthermore, two structural arrangements were tested: the IP positioned at the N-terminus of the shuttle (IP–linker–BBB shuttle) or at the C-terminus (BBB shuttle–linker–IP), to evaluate whether cargo position affects BBB homing and penetration.

The ability of the Cy7-labelled bi-functional peptides to cross the BBB was evaluated by fluorescent intravital imaging at 4 different timepoints following retro-orbital vein injection in wild type (wt) CD-1 mice. Fluorescence was observed in the brain 1h after injection for peptide PEP 1 (positive control) and PEP 4 but disappeared thereafter, likely due to rapid clearance (Figure 2). PEP 3 also showed brain fluorescence 1h after injection but this signal remained stable for at least 6h and disappeared only 24h post-retro-orbital vein injection (Figure 2A), suggesting slower elimination compared to peptides PEP 1 and PEP 4. The intensity of fluorescence in counts per second (CPS) is shown for PEP 3. We did not detect fluorescence in the brain upon injection of peptides PEP 2, PEP 5, PEP 6, PEP 7, PEP 8, PEP 9 and PEP 10. Figure 2B shows fluorescence for peptides PEP 5, PEP 7 and PEP 10. It is interesting to note that in peptides PEP 7 to PEP 10, the BBB shuttle is positioned at the N-terminus of the IP-LRRK2, suggesting this conformation disables BBB crossing.

**Figure 2.**
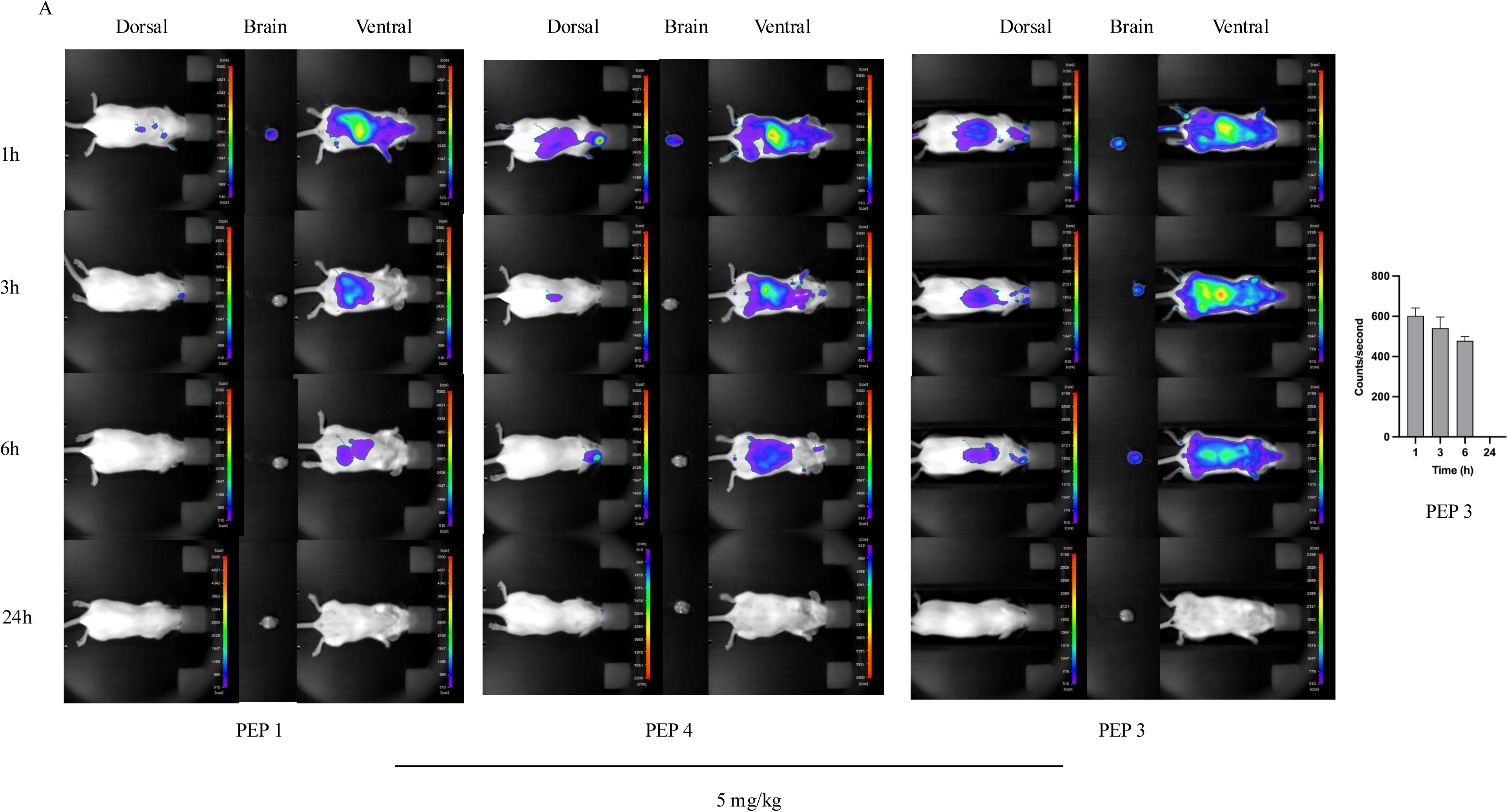
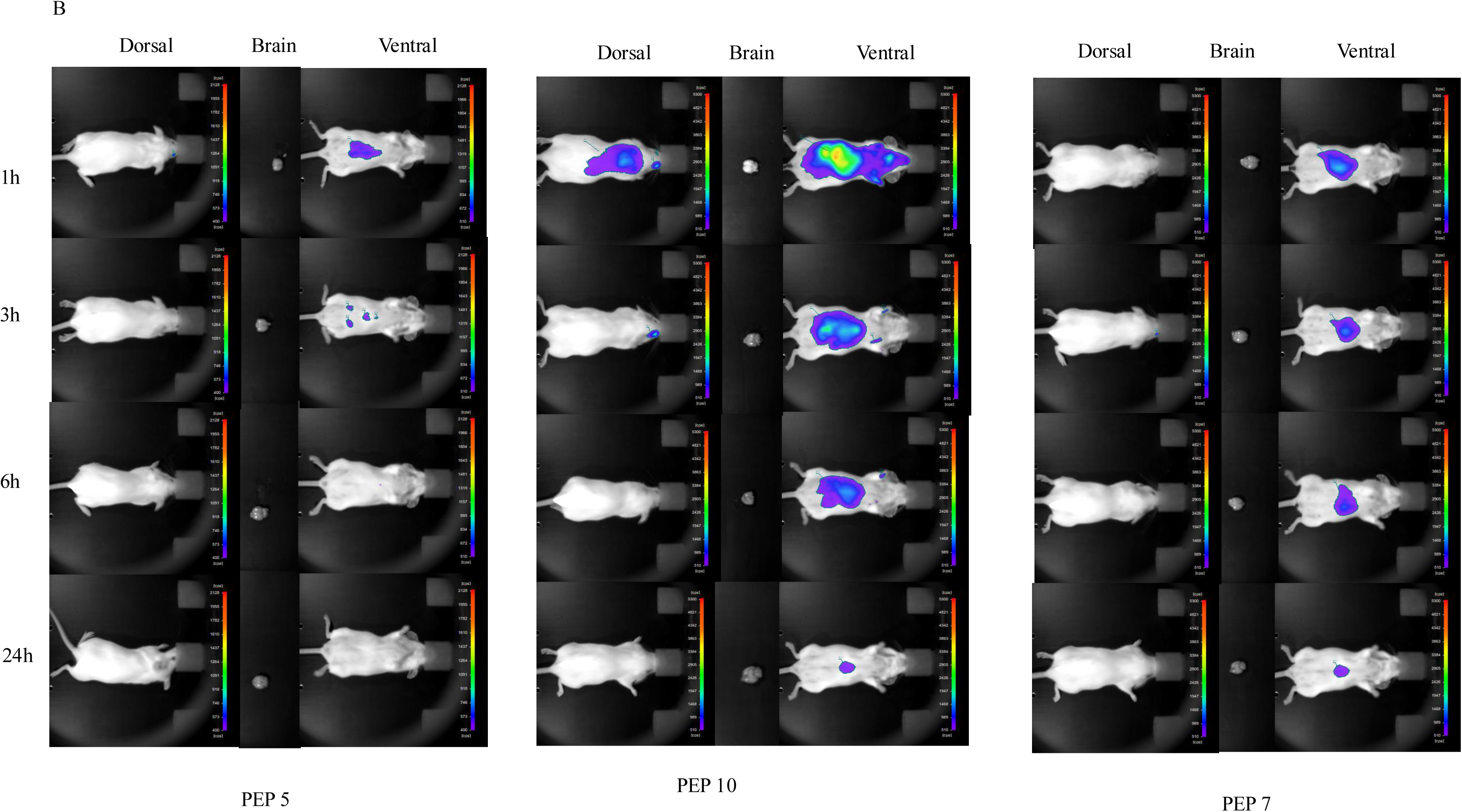
Fluorescent whole-body imaging showing the biodistribution of Cy7-labelled peptides PEP 1, PEP 3, PEP 4, PEP 5, PEP 7 and PEP 10. A) Mice (n=3) were injected with the Cy7-labelled PEP 1, PEP 3 and PEP 4 peptides (5 mg/kg) in the retro-orbital vein and images were acquired at 1, 3, 6 and 24h. The scale bar on the right shows fluorescence intensity where red represents highest intensity and purple lowest intensity. Scans show the dorsal and ventral positions, as well as the brain. Intensity of fluorescence in counts per second (CPS) is shown. B) Mice (n=3) were injected with Cy7-labelled PEP 5, PEP 7 and PEP 10 (5 mg/kg) in the retro-orbital vein and images acquired as in A.

Given that fluorescence of 5 mg/kg of PEP 3 bi-functional peptide persisted longer in the brain, we explored whether the intensity of the fluorescence was dose-dependent. Figure 3A shows images acquired 1h after retro-orbital vein injection using three different concentrations of Cy7-labelled PEP 3 (2.5, 5 and 10 mg/kg). The results indicate a clear dose-dependent relationship with higher doses resulting in higher fluorescence intensity in the brain. No detectable fluorescence was observed at the lowest tested dose of the peptide (2.5 mg/kg). The intensity of fluorescence in counts per second I (CPS) s shown.

**Figure 3.**
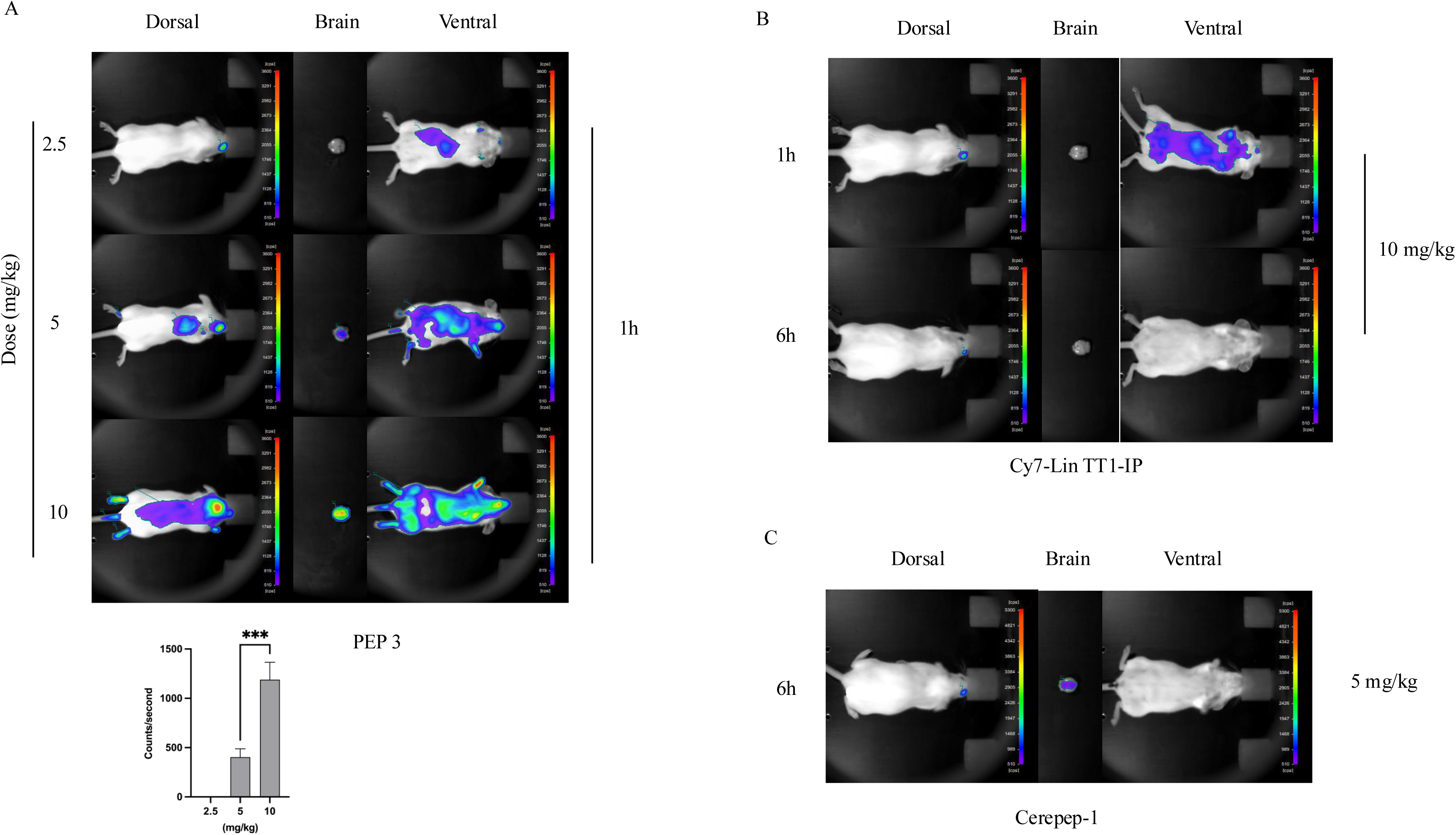
Fluorescent whole-body imaging showing the dose effect of the Cy7-labelled peptide PEP 3, Cy7 Lin TT1-IP and Cerepep 1. **A)** Mice (n=3 were injected in the retro-orbital vein with peptide Cy7-labelled PEP 3 at concentrations of 2.5, 5 and 10 mg/kg) and imaged 1h after injection. Dorsal and ventral positions, as well as the brain are shown. The scale bar on the right shows fluorescence intensity. **B)** The Cy7-labelled Lin TT1-IP peptide was injected in the retro-orbital vein at 10 mg/kg and mice (n=3) were imaged 1 and 6h after the injection. Dorsal and ventral positions, as well as the brain are shown. The scale bar represents fluorescence intensity. **C**) The Cy7-labelled Cerepep 1 shuttle was injected in the retro-orbital vein (5 mg/kg) and mice (n=3) were imaged 1h after the injection. Dorsal and ventral positions, as well as the brain are shown.

As a negative control, we used a high concentration of C7-labelled Lin TT1-IP, a peptide consisting of the p32 targeting tumor-penetrating peptide Lin TT1 fused to an IP. As expected, the Cy7-labelled peptide was not detected in the brain (Fig 3B). As a positive control, mice were injected with 5 mg/kg of the Cererpep-1 shuttle peptide, that yielded a fluorescence signal in the brain 1h after injection as expected (Fig 3C). As a control, peptide Lin TT1-IP is degraded by proteases recovering upon 24h of incubation in serum around 30% of the initial peptide (Fig 4B).

**Figure 4.**
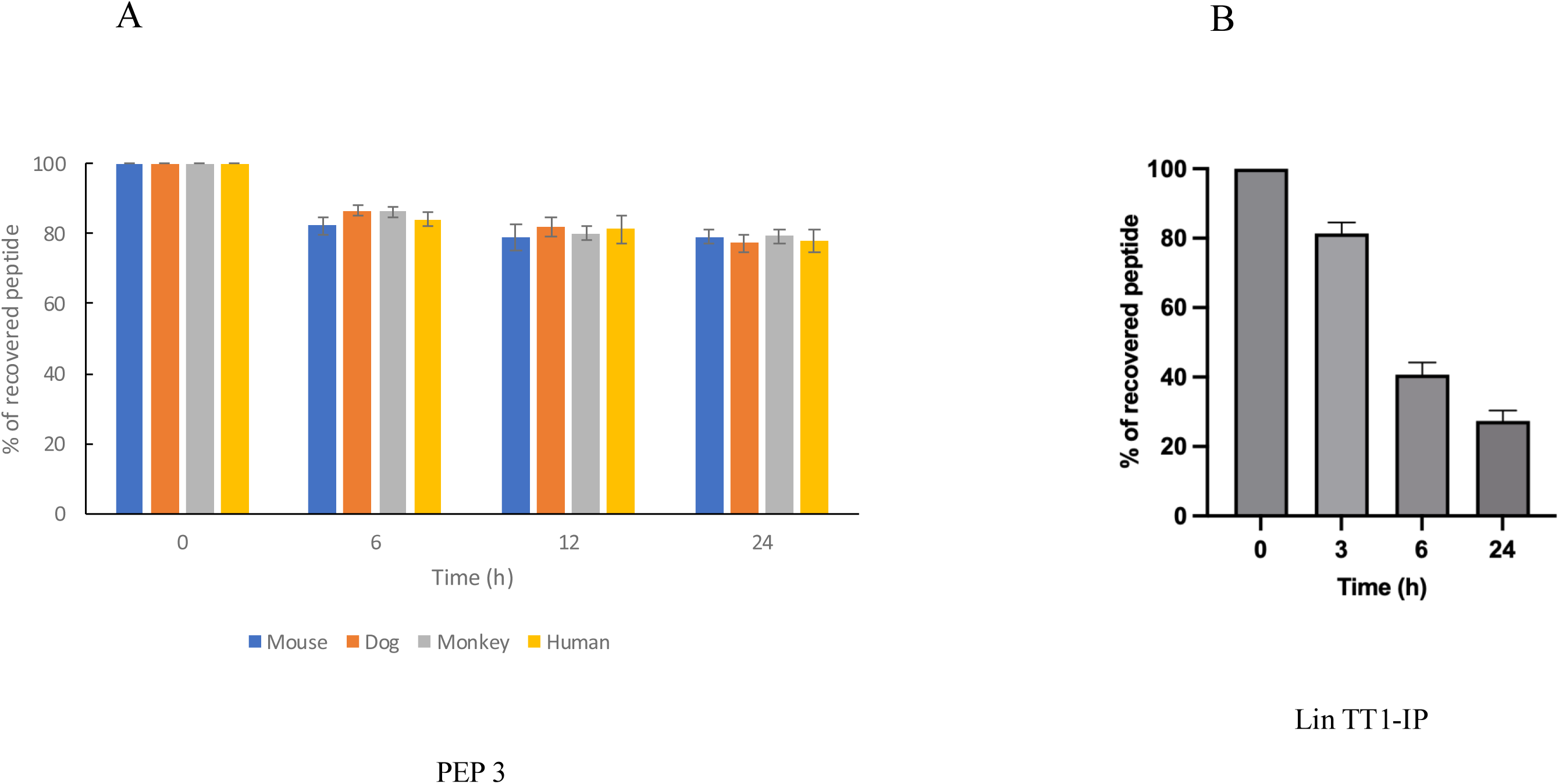
Stability of the PEP 3 peptide in mouse, human, dog and monkey serum. The PEP 3 peptide (A) was incubated at 37^°^C in serum of different species for several intervals from 0 to 24h. The Lin TT1-IP (B) peptide was incubated at 37°C with human serum from 1 to 24h.The integrity of the peptides was analysed by MS. The percentage of recovered peptide is represented. Every measurement was performed in triplicate. Similar results were obtained in two independent experiments. Standard deviation is shown.

### PEP 3 bi-functional peptide is stable to degradation by serum proteases

Proteolytic degradation of peptide-based drugs is a major drawback that often limits systemic therapeutic applications. We analyzed the stability of PEP3 peptide at 6, 12 and 24h after incubation at 37^°^C with human, mouse, dog and monkey serum by MS. Figure 4A shows that regardless of the species, after a slight drop between 0 and 24h post-incubation, 80% of the intact PEP 3 peptide is recovered, highlighting its stability. Thus, PEP 3 seems strongly resistant to degradation by serum proteases, making it a strong candidate for therapeutic development.

### Systemic PEP 3 treatment does not cause adverse effects in wild type mice

To assess the safety of PEP 3 bi-functional peptide, we recorded the body weight of 6 mice following a two weeks treatment, five days per week at a dose of 5 mg/kg i.p. As shown in Figure 5, PEP 3-treated CD1 wt mice showed no significant reduction in body weight, compared to the saline-treated control group. These findings suggest that systemic administration of PEP3 is well tolerated and does not cause adverse effects in wt mice.

**Figure 5.**
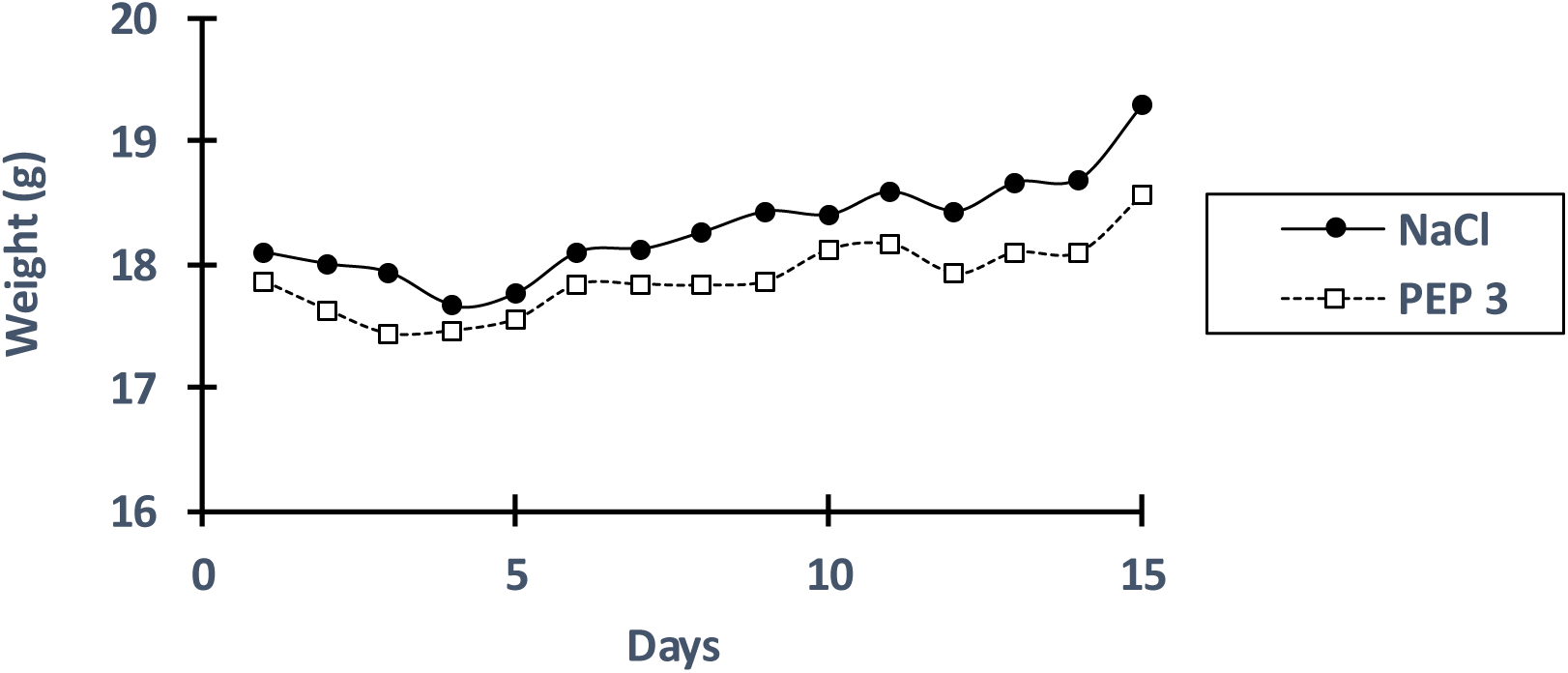
Effect of the PEP 3 peptide on body weight. PEP 3 peptide (5 mg/kg) or PBS were injected i.p. to CD1 mice (n=6/group) for 2 weeks and body weight (g) was monitored daily.

### Validation of the safety and therapeutic efficacy of PEP 3 peptide in a mouse model of PD

A model of PD was developed in CD-1 adult male mice by overexpressing alpha-synuclein locally and bilaterally in the SN. Two months after surgery, one cohort (n=20) was treated for 2 w and the other cohort (n=20) for 4 w with a single dose of PEP 3 peptide (7.5mg/kg, i.p.) or PBS (i.p.).

Overall, no adverse events were recorded after intracerebral injections of the AAV virus, the generation of the model or the repeated i.p. injections. No changes in fur appearance, grooming behavior, food intake or social interactions were observed, and body weight remained stable throughout in all cohorts, suggesting the PEP 3 treatment was well tolerated. A brief freezing of behavior was noticed immediately (>2 minutes) following i.p. administration of PEP 3 without further physiological or behavioral consequences observed. The 4 weeks treatment cohort presented hardenings at the i.p. injection site. It is worth noting, however, that a different administration route than repeated localized i.p. injection will be used in the clinic.

The therapeutic efficacy of PEP 3 was evaluated by immunohistochemical analysis of dopaminergic integrity and the levels of pathological phosphorylated alpha-synuclein. After two weeks of PEP 3 treatment, tyrosine hydroxylase (TH) staining in the SN of mice was markedly preserved, compared to PBS-injected control parkinsonian mice (p=0.001) (Figure 6 A-E). The same cohort showed a reduction of pathological alpha-synuclein (P129) in the SN (p=0.034; Figure 7 A-F), suggesting that PEP 3 protects dopaminergic neurons from degenerating and reduces alpha-synuclein phosphorylation in this mouse model of PD.

**Figure 6.**
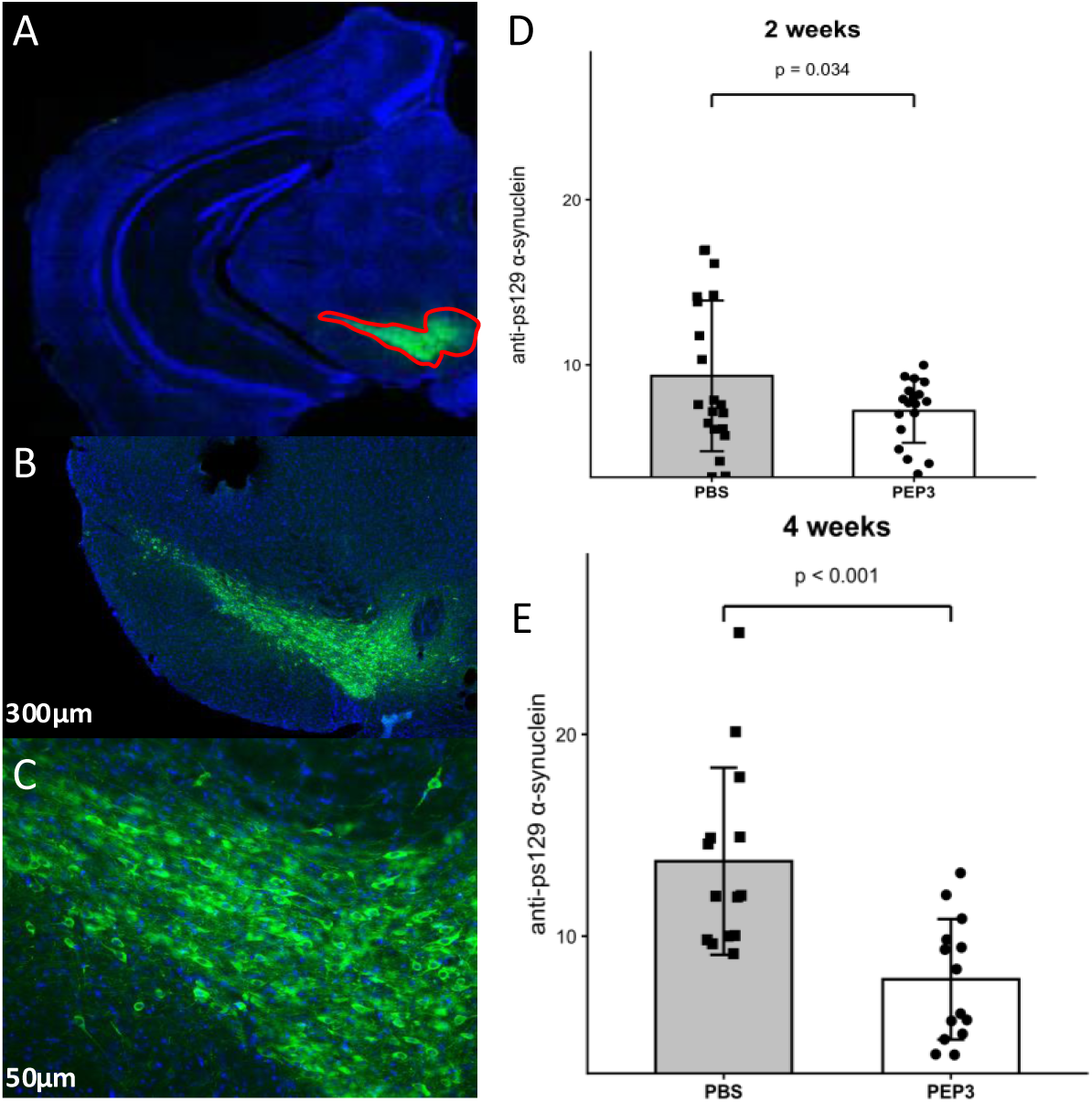
Effect of the PEP 3 peptide on TH fluorescence. Coronal sections (n=12-15/mouse) stained with TH showing labelled neurons in the SN (green) in 2 w PEP3-treated mice (n=10/group) (A, B, C). Quantification of TH fluorescence intensity in PEP 3-treated and PBS control groups at 2 w (D; p= 0.001, g=1.05) and 4 w (E; p=0.017, g=0.70).

**Figure 7.**
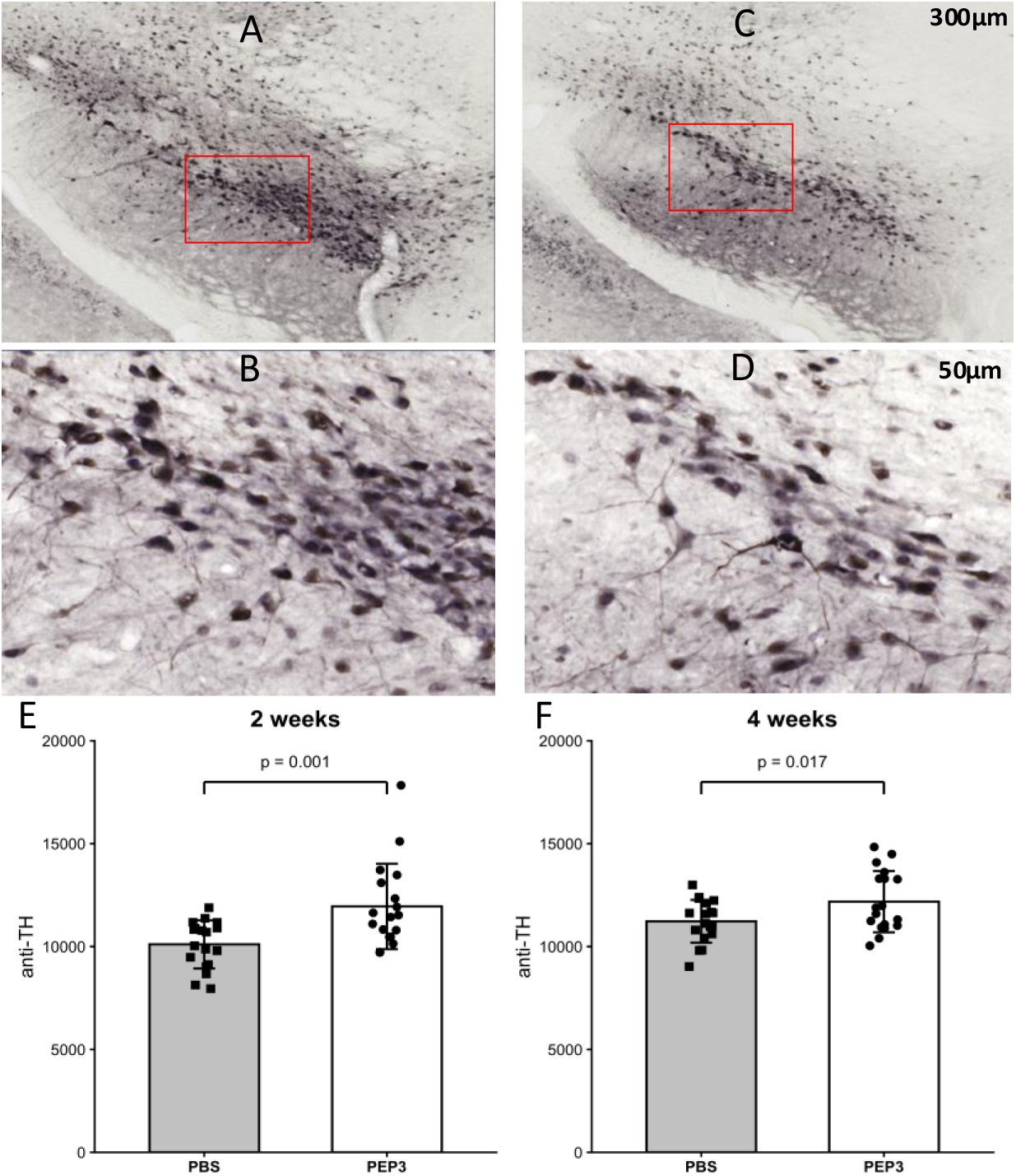
Effect of the PEP 3 peptide on alpha-synuclein staining. Coronal sections (n=12/mouse) stained with an anti-PS129-alpha-synuclein antibody showing labelled neurons in the SN in 4 w PEP 3-treated and control mice (n=10/group) (A, B, C, D). Quantification of PS129-alpha-synuclein staining expressed as a percentage of the surface of the region of interest at 2 w (E; p=0.034, g=0.57) and 4 w (F; p<0.001, g=1.05) post-treatment.

To further validate these findings, a second cohort of parkinsonian mice was treated for 4 weeks i.p. Consistent with the results obtained after 2 w of treatment, the post-mortem analysis confirmed a significant dopaminergic neuron integrity (TH, p=0.017). Moreover, a very large reduction in phosphorylated alpha-synuclein (p<0.001) was observed (Figure 7E). These data suggest that PEP3 interference with LRRK2 contributes to the decrease in alpha-synuclein phosphorylation and resulting pathology. Importantly, the efficacy observed in both cohorts also confirms that PEP 3 crosses the BBB in a pathological PD model.

## DISCUSSION

Parkinson’s disease (PD) is the second most common neurodegenerative disorder worldwide and is characterized by motor and cognitive deficits resulting from the progressive loss of dopaminergic neurons, neuroinflammation and alterations in the neurotransmitter system. The degeneration of dopaminergic neurons is accompanied by the accumulation of pathological alpha-synuclein in specific brain regions, which evolves over time and contributes to the broad clinical heterogeneity observed in PD patients (***4, 25–27***).

Protein/protein interactions (PPIs) play a central role in the regulation of cellular homeostasis, and their dysregulation is implicated in numerous diseases, including cancer and neurodegenerative disorders(***16***). Targeting PPIs therefore represents an attractive therapeutic strategy. In the present study, we focused on modulating the interaction between the phosphatase (PP1) and leucine-rich repeat kinase (LRRK2), a kinase strongly associated with familial and sporadic forms of PD. In fact, impaired alpha-synuclein clearance is a central feature of PD pathology, leading to its accumulation and aggregation. Genetic factors, including mutations in LRRK2, have been shown to interfere with normal alpha-synuclein turnover, while alterations in enzymes involved in neurotransmitter metabolism may further exacerbate this process (***26, 28, 29***). Phosphorylation is the most prevalent post-translational modification of _α_-synuclein. Although _α_-synuclein phosphorylation occurs at low levels under physiological conditions, pathological states are associated with increased phosphorylation at specific serine residues. In particular, phosphorylation at serine 129 has been detected in Lewy bodies from PD patients and is strongly implicated in disease progression (***17, 30***).

We identified a 13-amino-acid region within the LRRK2 protein that adopts an _α_-helical conformation and mediates its interaction with PP1 (***23***). The ability of the corresponding interfering peptide (IP-LRRK2) to disrupt this interaction was confirmed using *in vitro* competition assays.

In order to enable central nervous delivery, IP-LRRK2 was conjugated to several BBB-penetrating peptide shuttles, generating bi-functional peptides that retained their capacity to disrupt the PP1/LRRK2 interaction. These constructs were subsequently evaluated in preclinical *in vivo* studies using well-established mouse models of PD.

LRRK2 is phosphorylated at multiple sites, and although the precise mechanisms governing its phosphorylation dynamics are not fully understood (***18, 21, 22, 31, 32***), alterations in its phosphorylation status are clearly linked to PD pathogenesis (***19***). In addition, PP1 has been identified as a major phosphatase responsible for LRRK2 dephosphorylation (***16***), highlighting the functional relevance of the PP1–LRRK2 interaction. The IP-LRRK2 peptide described here therefore represents a valuable tool to modulate the phosphorylation of LRRK2 indirectly by controlling PP1 binding, offering a novel approach to influence LRRK2-dependent signaling pathways in PD.

The therapeutic efficacy of the newly generated bi-functional peptides was evaluated preclinically in a well-characterized mouse model of PD (***28, 29, 33***). PEP 3 was shown to reduce levels of phosphorylated-α-synuclein, a key driver of PD pathology. In addition, PEP 3 treatment protected the dopaminergic neurons, as evidenced by the higher levels of TH staining in the striatum of parkinsonian treated mice compared to untreated controls. These findings strongly support the further preclinical development of PEP3, including studies in large animal models to evaluate clinically relevant routes of administration and generate predictive data on pharmacokinetics, biodistribution, target engagement, and preliminary safety.

The BBB plays a crucial role in the maintaining homeostasis in the central nervous system by restricting the entry of potential harmful substances (***34***). However, this protective function also limits the effectiveness of many therapeutics for neurological diseases. Peptide-based BBB shuttles have emerged as a promising strategy to overcome this limitation by enabling non-invasive delivery of bioactive molecules to the brain (36–38). Moreover, peptide shuttles offer several advantages over alternative brain-delivery approaches, including low toxicity, reduced immunogenicity, absence of drug resistance, and straightforward chemical synthesis. The BBB shuttle peptides employed in this study successfully facilitated the transport of an interfering peptide into the brain, supporting their utility as a delivery platform for PD therapeutics. Continued advances in BBB shuttle design are expected to accelerate preclinical development and clinical translation of peptide-based therapies for PD.

## CONFLIC OF INTEREST

Authors declare no conflict of interest

